# An interolog-based barley interactome as an integration framework for immune signaling

**DOI:** 10.1101/2021.11.02.466982

**Authors:** Valeria Velásquez-Zapata, J. Mitch Elmore, Greg Fuerst, Roger P. Wise

## Abstract

The barley MLA nucleotide-binding, leucine-rich-repeat (NLR) receptor and its orthologs confer recognition specificity to many cereal diseases, including powdery mildew, stem and stripe rust, Victoria blight, and rice blast. We used interolog inference to construct a barley protein interactome (HvInt) comprising 66133 edges and 7181 nodes, as a foundation to explore signaling networks associated with MLA. HvInt was compared to the experimentally validated Arabidopsis interactome of 11253 proteins and 73960 interactions, verifying that the two networks share scale-free properties, including a power-law distribution and small-world network. Then, by successive layering of defense-specific ‘omics’ datasets, HvInt was customized to model cellular response to powdery mildew infection. Integration of HvInt with expression quantitative trait loci (eQTL) enabled us to infer disease modules and responses associated with fungal penetration and haustorial development. Next, using HvInt and an infection-time-course transcriptome, we assembled resistant (R) and susceptible (S) subnetworks. The resulting differentially co-expressed (R-S) interactome is essential to barley immunity, facilitates the flow of signaling pathways and is linked to *Mla* through trans eQTL associations. Lastly, next-generation, yeast-two-hybrid screens identified fifteen novel MLA interactors, which were incorporated into HvInt, to predict receptor localization, and signaling response. These results link genomic, transcriptomic, and physical interactions during MLA-specified immunity.

**AUTHOR SUMMARY:** Powdery mildew fungi infect more than 9,500 agronomic and horticultural plant species. In order to prevent economic loss due to diseases caused by pathogens, plant breeders incorporate resistance genes into varieties that are grown for food, feed, fuel and fiber. One of these resistance genes encodes the barley MLA immune receptor, an ancestral cereal protein that confers recognition to powdery mildew, stem and stripe rust, rice blast and Victoria blight. However, in order to function properly, these immune receptors must interact with additional proteins and protein complexes during the different stages of fungal infection and plant defense. We used a combination of computational- and laboratory-based methods to predict over 66,000 possible protein-protein interactions in barley. This network of proteins was then integrated with various defense-specific datasets to assemble the molecular building blocks associated with resistance to the powdery mildew pathogen, in addition to those proteins that interact with the MLA immune receptor. Our application of genome-scale, protein-protein interaction data provides a foundation to decipher the complex molecular components that control immune responses in crops.

## INTRODUCTION

Plants attain a focused immune response via interconnected signaling networks (Yuan *et al*., 2020; Ngou *et al*., 2021). These networks are activated upon recognition of conserved pathogen molecules termed pathogen-associated molecular patterns (PAMPS) (Lo-Presti *et al*., 2015; Toruño *et al*., 2016; Bentham *et al*., 2020). Pathogens have adapted to suppress this response by secreting effectors into the host cell. As a counter mechanism, families of resistance (R) proteins have evolved, encoding a series of intracellular receptors that are associated with immune activation (Lo-Presti *et al*., 2015; Toruño *et al*., 2016). Experimental and computational evidence have shown that most R-proteins converge to a group of intracellular nucleotide-binding, leucine-rich-repeat (NLR) receptors, one of the largest plant protein families (Van-Wersch *et al*., 2020; Monteiro and Nishimura, 2018; Baggs *et al*., 2017; Sun *et al*., 2020; Tamborski and Krasileva, 2020). The protein structure of this family consists of an N-terminal domain (coil-coiled, CC, or Toll-like Interleukin-1 receptor Resistance protein, TIR), a conserved nucleotide-binding domain shared with APAF-1, various R-proteins and CED-4 (NB-ARC), and a C-terminal leucine-rich repeat domain (LRR) (Van-Wersch *et al*., 2020; Monteiro and Nishimura, 2018; Baggs *et al*., 2017). Since R-proteins have a prevalent role in determining the outcome of plant-pathogen interactions, they are ideal probes to dissect host immunity as well as basic plant cell function (Bray Speth *et al*., 2007; Sun *et al*., 2020).

Obligate biotrophic fungi, which include mildews and rusts, cause some of the most detrimental impacts to crop production (Dean *et al*., 2012). In the interaction between *Blumeria graminis* f. sp. *hordei* (*Bgh*), the causal agent of powdery mildew, and its cereal host plant, barley (*Hordeum vulgare* L.), disease is blocked by the action of specific barley R-proteins, designated ML, that respond to corresponding *Bgh* effectors, designated AVR (Lu *et al*., 2016; Saur *et al*., 2019; Bauer *et al*., 2021; Ridout *et al*., 2006). Diversification of the *Mla* NLR (Wei *et al*., 2002; Seeholzer *et al*., 2010), has generated up to 30 allelic variants that confer specific recognition to different AVR-associated *Bgh* isolates (Lu *et al*., 2016). MLA-mediated signaling involves the stabilization of the protein through interactions with the HRS complex, which consists of three co-chaperones: the heat shock protein 90 (HSP90), the Zn^2+^ binding protein Required for Mla12 Resistance 1 (RAR1), and the suppressor of G-two allele of Skp1 (SGT1) with direct binding with MLA for the first two (Bieri *et al*., 2004; Shirasu, 2009). A RING-type E3 ligase (MIR1) has also been reported as an interactor of MLA, attenuating defense signaling by degradation of the receptor via the ubiquitin proteasome system (Wang *et al*., 2016). Upon recognition of *Bgh* AVR effectors, MLA accumulates in the nucleus, followed by its association with multiple transcription factors (TFs), including WRKY1, WRKY2 and MYB6 (Chang *et al*., 2013; Shen *et al*., 2007).

In order to establish a comprehensive view of the regulatory programs that render a plant resistant to pathogens, deeper insight into *R*-gene activation and the resulting signaling cascades is needed (Deng *et al*., 2020; Martin *et al*., 2003). In this regard, protein interactomes are advantageous to interpret the molecular mechanisms in immune cell signaling (Mukhtar *et al*., 2011; Weßling *et al*., 2014). Yet, building interactomes from experimental data is a cost and labor intensive task, and as a result, they are incomplete and only available for few model organisms (Matthews *et al*., 2001; McWhite *et al*., 2020). To circumvent this challenge, predictive approaches have been developed. One of these is interologs inference, which consists of mining protein interactions using ortholog information between the species of interest and model organisms that possess experimentally validated interactions (Matthews *et al*., 2001). This approach has been successfully used to generate predicted interactomes in Arabidopsis and several crop species including rice, maize and tulsi (Musungu *et al*., 2015; Ho *et al*., 2012; Geisler-Lee *et al*., 2007; Singh *et al*., 2020). Then, integration of interactomes with context-specific data types can be performed to highlight nodes, edges, structures, and modules associated with a phenotype of interest (Randhawa and Pathania, 2020). For example, infection time-course transcriptomes enable the use of gene co-expression as evidence to increase confidence of predicted protein-protein interactions (PPIs) and build phenotype-specific subnetworks (Braun *et al*., 2013; Petrey and Honig, 2014; Jiang *et al*., 2016). In addition, eQTL data can be used to obtain disease modules in the interactome and provide genetic and physical connections between nodes, avoiding biases given by the amount of information of well-studied disease genes (Wang *et al*., 2020; Dwivedi *et al*., 2020).

To discover new connections in NLR-based immunity, we used interologs inference to develop a predicted interactome for the Triticeae grain crop, barley, using experimentally validated interactions in several model organisms and crop species. Then, using host-pathogen-associated gene expression and eQTL data, we assembled several subnetworks associated with resistance, susceptibility, and the dynamics of defense during key stages in powdery mildew infection. Lastly, we used yeast-two-hybrid, next-generation-interaction screening (Y2H-NGIS) to identify and validate new protein-protein interactions (PPIs) between the MLA6 NLR-receptor and barley targets and extended this to additional MLA alleles. Fifteen novel interactors were identified and overlayed onto the predicted barley interactome and by extension, cellular compartments. These analyses enabled us to propose a model of MLA cellular localization across *Bgh* infection and to associate the receptor with gene co-expression during resistance.

## RESULTS

### The barley predicted interactome shows protein essentiality in defense

We generated a high-quality, barley (*H. vulg*are) interactome (HvInt) based on interologs (Matthews *et al*., 2001). HvInt contains 66133 edges and 7181 nodes (S1 Data) and can be used as baseline network to investigate different signaling events in the barley cell. Here we show how HvInt can be customized to study immune response, via integration of defense-specific datasets. Figure 1A illustrates the number of interactions mined from the different species used for interologs prediction. About 87.8% of the predicted interactions are supported by experimental validations in *Arabidopsis thaliana* (including the compiled interactions in the pan-plant protein interactome (McWhite *et al*., 2020)), followed by *Saccharomyces cerevisiae* (11.8%), *Oryza sativa* (0.012%) and *Zea mays* (0.0005%). Properties of HvInt were compared to the collected experimentally validated interactions in *A. thaliana* (AtInt), which comprise 73960 interactions and 11253 proteins (S1 Data). This analysis demonstrated that HvInt maintains the power-law and small-world properties of the AtInt network (p-value > 0.05 for the kolmogorov-smirnov tests, global clustering coefficient ratio >1 and an average shortest path length ratio ∼ 1) (Albert and Jeong, 1999; Watts and Strogatz, 1998).

**Figure 1.**
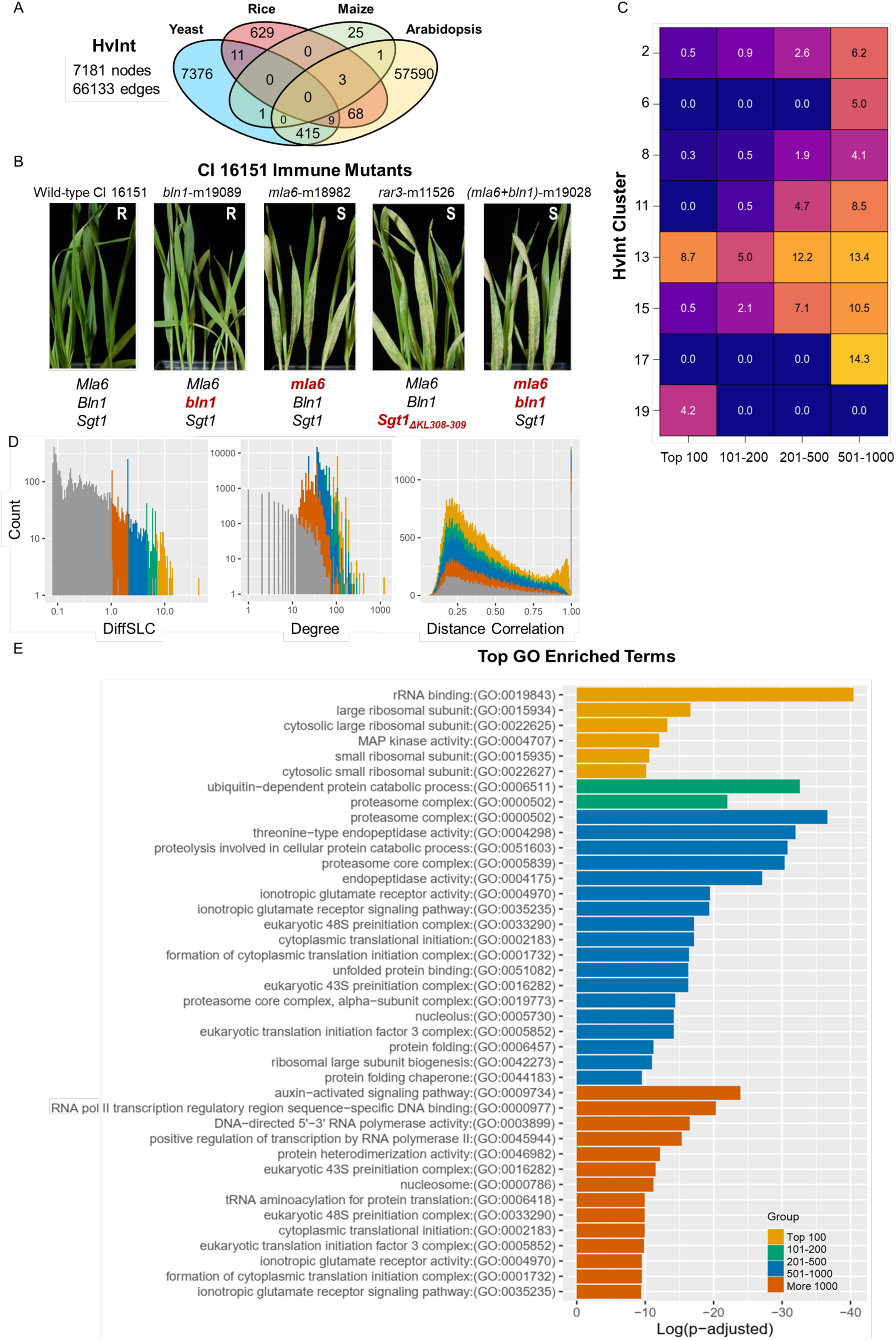
HvInt, the predicted barley interactome. A) Venn diagram with the number of interactions in HvInt across the species of origin. B) Wild-type CI 16151 (*Mla6, Bln1, Sgt1*) and derived immune mutants *bln1*-m19089 (*Mla6, bln1, Sgt1*), *mla6*-m18982 (*mla6, Bln1, Sgt1), rar3-*m11526 (*Mla6, Bln1, Sgt1*_*ΔKL308-309*_, and *(mla6+bln1)*-m19028 (*mla6, bln1, Sgt1)* 7 days after inoculation with *Bgh* isolate 5874 (*AVR*_*a6*_). R = resistant, S = susceptible; mutant alleles are designated below the images in red. C) Distribution of the top essential proteins in HvInt across clusters. D) HvInt properties associated with the top essential proteins. E) GO term enrichment of the top essential proteins in HvInt.

We applied a walktrap clustering algorithm on HvInt using distance correlation from expression data as edge weights (Székely *et al*., 2007). For this purpose we used RNA-Seq expression data from an infection time course with *Bgh* isolate 5874 (*AVR*_*a6*_) [0, 16, 20, 24, 32 and 48 hours after inoculation (HAI)] of the resistant barley line CI 16151 (*Mla6, Bln1, Sgt1*) and four fast-neutron derived mutants; one resistant -- *bln1*-m19089 (*Mla6, bln1, Sgt1*), and three susceptible -- *mla6*-m18982 (*mla6, Bln1, Sgt1), rar3-*m11526 *(Mla6, Bln1, Sgt1*_*ΔKL308-309*_), and the double mutant m19028 (*mla6, bln1, Sgt1)*, as shown in Figure 1B (Chapman *et al*., 2020; Hunt *et al*., 2019). We acquired fifty-four clusters comprising 1 to 2313 proteins, with 97.6% in the top five clusters (2, 8, 11, 13 and 15). We then calculated node essentiality using DiffSLC (S2 Data), which correlates with functionally important nodes, whose deletion is associated with lethality (Mistry *et al*., 2017). The top 1000 essential proteins were separated into four groups including the top 100, 101-200, 201-500 and 501-1000. These essential proteins were distributed in eight clusters, including the top five most populated, as shown in Figure 1C, with significant enrichment in the clusters 13 and 15 (S2 Data). DiffSLC was compared to the main properties that are used for its calculation: the expression distance correlation and node degree (Figure 1D). We found that DiffSLC and degree were correlated; this clarifies, why the top 1000 essential proteins (13.9% of the nodes) are involved in 80.7% of the interactions in HvInt. This led us to conclude that essential proteins comprise a set of hubs in HvInt and that DiffSLC is a useful node property that can be used to characterize HvInt subnetworks.

Essential proteins were analyzed using Gene Ontology (GO) terms (Figure 1E). The top 100 were enriched with terms associated with ribosome and MAP kinase activity; 101-200 were associated with channel activity, rRNA binding, proteasome complex and ubiquitin-dependent catabolism; and 201-500 were associated with chaperone-mediated protein folding, auxin-activated signaling pathway, ionotropic glutamate receptor activity and endopeptidase activity. Finally, 501-1000 were associated with proton-exporting ATPase activity and translation. These results led us to conclude that some of the top essential proteins are linked with fundamental cellular processes while others have functions associated with plant immunity (Zeng *et al*., 2006; Forde and Roberts, 2014; Lee *et al*., 2015). Key examples are described in subsequent sections.

### Disease modules reveal barley cellular responses at *Bgh* penetration and haustorial development

Proteins associated with disease phenotypes tend to be tightly connected at the physical level, generating interaction modules and pathways (Barabási *et al*., 2011; Sharma *et al*., 2014). The concept of disease module emerges then to describe this phenomenon, comprising interactome clusters that are functionally linked to a disease phenotype through genetic and physical associations (Sharma *et al*., 2014). We adapted the disease-module concept, originally developed in the context of network medicine (Barabási *et al*., 2011), to add temporal-defense information to HvInt at two key infection stages by *Bgh*. To identify disease modules during *Bgh* infection, we used N2V-HC (Wang *et al*., 2020) to integrate HvInt with eQTL data derived from transcriptome analysis of the barley Q21861 × SM89010 doubled-haploid population during appressorial penetration (16 HAI) and haustorial development (32 HAI) (Surana *et al*., 2017). All genetic markers with significant eQTL associations (adjusted p-values < 0.001) were used as input, containing 317 markers mapped to 1009 genes and associated with 16943 eQTLs. From those, we filtered out associations with adjusted p-values larger than 0.001, ending up with 4357 eQTLs at 16 HAI and 6375 at 32 HAI. This resulted in disease modules with proteins enriched with eQTLs by timepoint (S3 Data).

We performed Gene Ontology (GO) analyses of the disease modules to further understand the temporal response to powdery mildew. Modules were associated with individual timepoints, and a core response common to both timepoints. At penetration, we found 584 significant GO terms (adjusted p-values < 0.05) from which 85 were unique to this developmental stage. Haustorial development had 773 significant enriched GO terms associated, 274 of them unique. The 499 GO terms associated with the core response, as well the 359 unique, were explored further by looking for enrichment of differentially expressed (DE) genes (adjusted p-values < 0.001) extracted from our RNA-Seq data described above. By comparing the resistant CI 16151 (*Mla6, Bln1, Sgt1)* and the derived susceptible mutant, m18982 (*mla6, Bln1, Sgt1)*, we could focus on gene sets that were functionally specific and displayed expression differences between compatible and incompatible interactions connected to the *Mla6* NLR. The core response consisted of 110 GO terms that had DE genes associated at 16 HAI and 32 HAI (shown in S1 Figure). The most significant were associated with vesicle trafficking, transporter activity, serine/threonine kinase signaling, and response to biotic stimulus, which is consistent for a host-pathogen interaction scenario. Unique GO terms at appressorial penetration included protein import into mitochondrial matrix and the polyamine biosynthetic process. In contrast, haustorial development was characterized by the unique GO terms glycine biosynthetic process from serine, cell wall macromolecule catabolic process, chitin catabolic process, carbon fixation, pyruvate metabolic process and carbohydrate phosphorylation.

Figure 2A provides an overview of the various DE genes associated with the core and unique GO terms by timepoint, which we designated disease module DE (DM DE) genes (S1 Figure, S3 Data). Our analyses indicate that these genes play a key role in the disease-module functions at each infection timepoint. The 16- and 32 HAI core response includes the DM DE genes: alcohol dehydrogenase, three kinases, receptor-like protein kinase, glutamate receptor and aquaporin. During fungal penetration (16 HAI), 30 unique DM DE genes, including two WRKY and one agamous MADS-box TF, two DETOXIFICATION proteins and several kinases were found. In contrast, during haustorial development (32 HAI) we identified 67 DM DE genes including chitinase, MLO-like protein, leucine-rich repeat receptor-like protein, malate dehydrogenase, proteasome-associated proteins, and several transporters. To interrogate activity over time, DM DE genes associated with the core and unique responses were visualized by expression profile plots for the CI 16151 (*Mla6, Bln1, Sgt1)* and m18982 (*mla6, Bln1, Sgt1)* genotypes (examples in Figure 2B). Expression profiles were characterized by significant differences between the two genotypes across all six timepoints, or by contrast, displayed an abrupt expression change at penetration, or haustorial development, in one of the genotypes.

**Figure 2.**
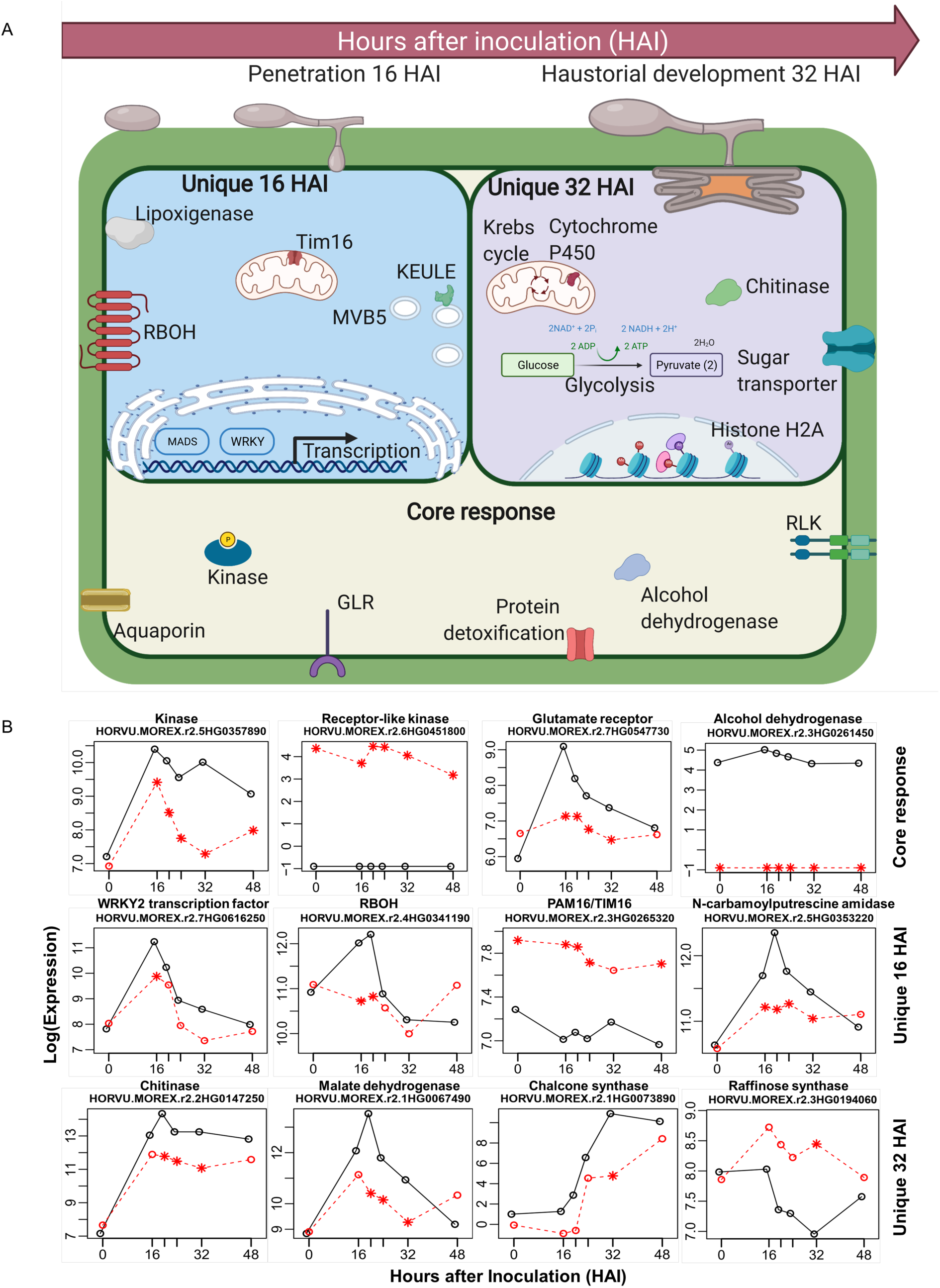
Identification of disease modules at *Bgh* appressorial penetration and haustorial development. A) Schematic summary of the DM DEs at penetration (16 HAI) and haustorial development (32 HAI), separated by core and unique responses. B) Examples of time-course expression patterns of DM DE genes by type of response, CI 16151 (*Mla6, Bln1, Sgt1)* in black and m18982 (*mla6, Bln1, Sgt1)* in red, significant differences (adjusted p-values <0.001) are designated by an asterisk.

### Construction of co-expressed interactomes during resistance and susceptibility

Time-course RNA-Seq data empowers rapid visualization of differential transcript accumulation in wild-type vs. mutant backgrounds, and thus, in what way immune signaling pathways can be impacted by pathogens. To complement the disease module analysis above, we identified immune-active sub-networks under resistant and susceptible disease outcomes by integrating HvInt with our infection-time-course RNA-Seq data (Hunt *et al*., 2019; Chapman *et al*., 2020). The RNA-Seq read counts for each group of infection phenotypes in Figure 1B was used to calculate co-expression values defined as pairwise distance correlations across genes. HvInt was then subset into the resistant [HvInt(R)] and susceptible [HvInt(S)] interactomes by retaining significantly co-expressed pairs under each phenotype. We defined significant correlation values for each subnetwork using a permutation test (Figure 3A, see Methods).

**Figure 3.**
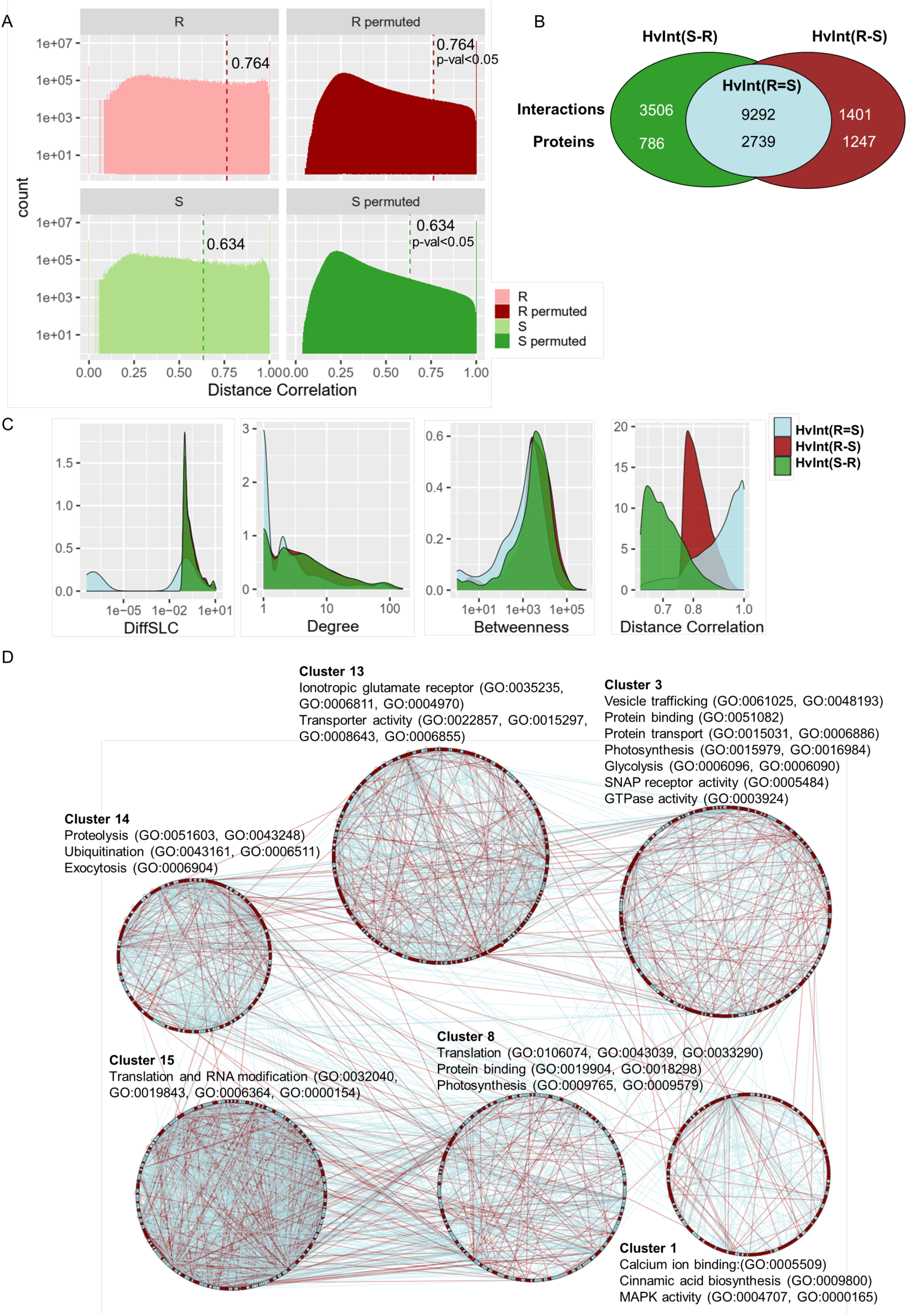
Construction of resistant and susceptible interactomes. A) Experimental and permuted distributions of the expression distance correlations in HvInt by disease phenotype, Resistant (R) or Susceptible (S); significance thresholds (p-value<0.05) are marked. B) Distribution of interactions and proteins in the resistant [HvInt(R)], susceptible [HvInt(S)] interactomes, separating by common [HvInt(R=S)] and difference [HvInt(R-S), HvInt(S-R)] subnetworks. C) Network properties of the resistant interactome, separated by common interactions with the susceptible network in blue [HvInt(R=S)] and the resistant unique interactions in red [HvInt(R-S)]. D) Clustering and GO annotation of the resistant interactome HvInt(R), separating the differentially co-expressed subnetwork in red HvInt(R-S) and the common HvInt(R=S) in light blue. Each cluster is grouped in a circular layout.

The resistant HvInt(R) subnetwork includes all edges with high expression distance correlation for the resistant genotypes. This subnetwork contains 3120 nodes and 10693 interactions (S4 Data) and was clustered into 33 groups with GO enrichment annotations as shown in S4 Data. The filtered susceptible subnetwork HvInt(S) has 12798 interactions and 3525 proteins, clustered in 45 modules with GO annotations reported in S4 Data. There is an overlap of 9292 interactions and 2739 proteins between the HvInt(R) and HvInt(S) subnetworks [Figure 3B, HvInt(R=S)]. We removed the common interactions HvInt(R=S) from HvInt(R) to obtain the differentially co-expressed resistant interactome HvInt(R-S) with 1401 interactions and 1247 proteins (S4 Data). We postulate that HvInt(R-S) shows unique, co-expressed interactions associated with the barley immune response to *Bgh*, therefore it represents the focus of our following analyses.

We tested for differences among the common HvInt(R=S), the differentially co-expressed resistant HvInt(R-S), and susceptible HvInt(S-R) interactomes by computing different topological properties that can be associated with the information content and flow in the networks. DiffSLC, degree, betweenness and expression distance correlation between the three subnetworks were compared as shown in Figure 3C. In addition, Wilcoxon rank sum tests were performed to quantify the significance of the differences. Results showed that HvInt(R-S) and HvInt(S-R) had significantly higher DiffSLC, degree and betweenness and lower expression distance correlation than HvInt(R=S) (p-values < 1×10^−100^). These results indicate that the proteins in the unique HvInt(R-S) and HvInt(S-R) have higher essentiality than the common resistant and susceptible HvInt(R=S) subnetwork. Importantly, these differences are associated with higher degree centrality, and not higher distance correlation, as the latter was lower in the differentially co-expressed subnetworks. We focused the following analyses on the resistant interactomes, as we were interested in exploring their functional significance.

The differentially co-expressed, resistant subnetwork HvInt(R-S) was analyzed using GO term enrichment. Figure 3D shows the HvInt(R), differentiating nodes and edges from HvInt(R-S) in red, and HvInt(R=S) in light blue. Community analysis revealed different distributions of these subnetworks in each cluster, retaining GO enriched terms that have been associated with plant immunity including: calcium ion binding, protein ubiquitination, and MAP kinase activity for the HvInt(R) cluster 1; vesicle trafficking, protein binding and transport, SNAP receptor activity and GTPase activity for HvInt(R) cluster 3; translation, protein binding and photosynthesis for HvInt(R) cluster 8; ionotropic glutamate receptor signaling pathway and transporter activity for HvInt(R) cluster 13. All terms are reported in S4 Data. Finally, using hypergeometric tests, enrichment of eQTL associations were calculated for each subnetwork, taking HvInt as reference. eQTLs were split by timepoint, using genome-wide or only *Mla* associations, a previously reported *trans* eQTL hotspot (Surana *et al*., 2017). From all the listed tests, significant enrichment (adjusted p-values <0.001) of *Mla* eQTL associations was only identified in the resistant networks HvInt(R) and HvInt(R-S) at 32 HAI (test adjusted p-values 7.76×10^−5^ and 5.56×10^−13^, respectively). From a total of 1247 proteins in HvInt(R-S), 299 are associated with the *Mla trans* eQTL, accounting for 23.9% of the total number of nodes (1.7 times more than HvInt which contains only 14% of associations).

### Novel interactors of MLA predict NLR localization and signaling

We used yeast-two-hybrid, next-generation-interaction-screening (Y2H-NGIS) to identify interacting partners for the MLA6 resistance protein, split into three baits corresponding to the main domains of the NLR: amino acids 1-161 for the coil-coiled (CC) domain, amino acids 1-225 for CC and the nucleotide binding (NB) domain (CC+NB), and amino acids 550-976 for the leucine rich repeat (LRR) domain (Velásquez-Zapata *et al*., 2021). Using the recently developed NGPINT (Banerjee *et al*., 2021) and Y2H-SCORES (Velásquez-Zapata *et al*., 2021) software, we obtained a list of high-confidence interactors for each of the MLA6 fragments and validated them using binary Y2H (Dreze *et al*., 2010). A total of three interactors were validated for the MLA6_CC_ domain and fifteen for MLA6_CC+NB_ domain (including the three interactors that interact with MLA6_CC_). No preys were validated for MLA6_LRR_. Table 1 shows the set of validated interactions for the MLA6 fragments, their descriptions, the *A. thaliana* orthologs, and the predicted cellular location in this organism in the TAIR database (Berardini *et al*., 2015). The confirmed interactors with MLA6_CC+NB_ had diverse predicted cellular localizations including nuclear, cytoplasmic, and organelle-associated proteins. Predicted functions of the MLA protein targets included transcription, vesicle transport, protein folding and degradation, among others. These predictions suggest novel localizations of the MLA receptor and molecular mechanisms that regulate its function in disease resistance.

**Table 1.**
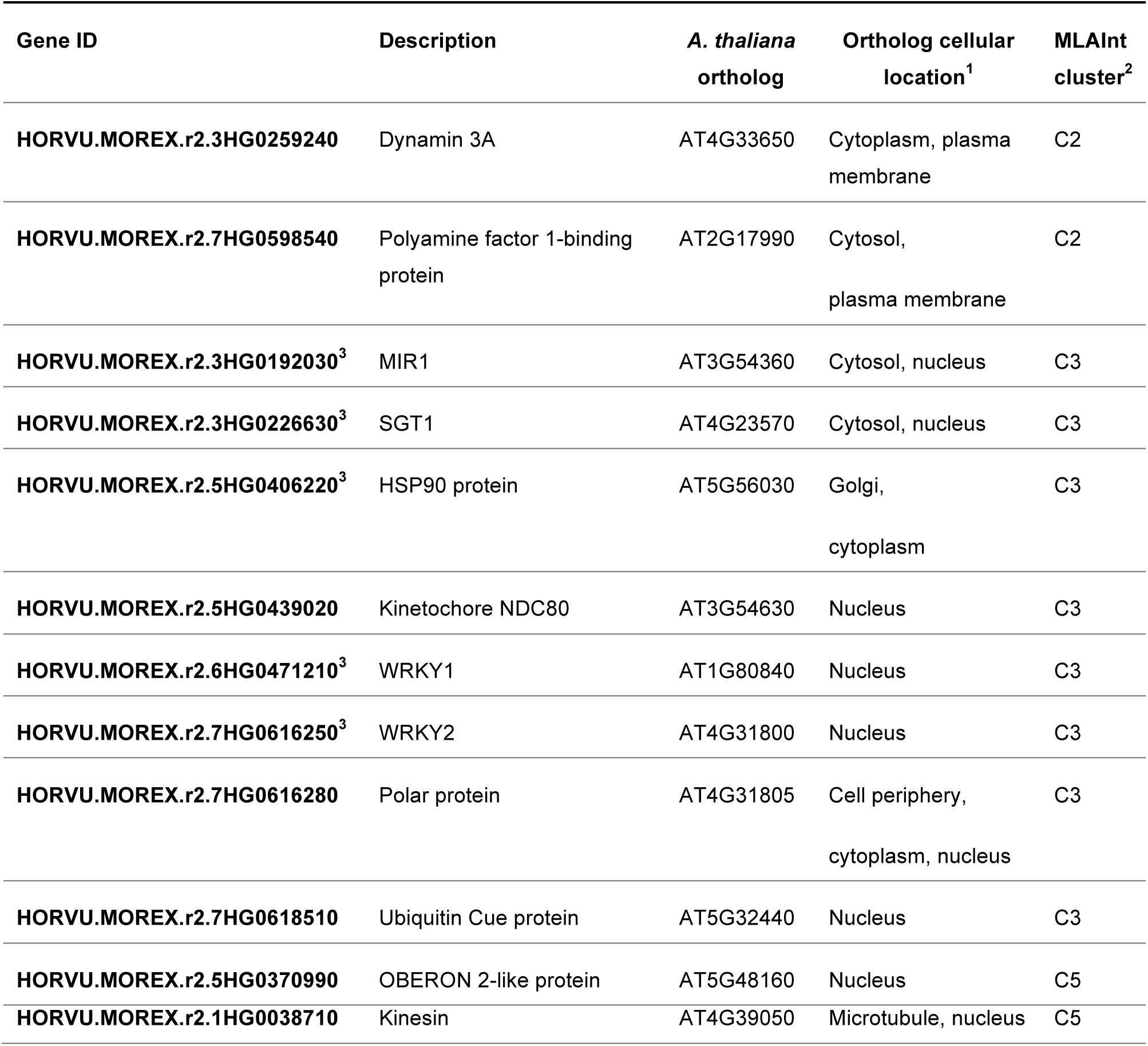
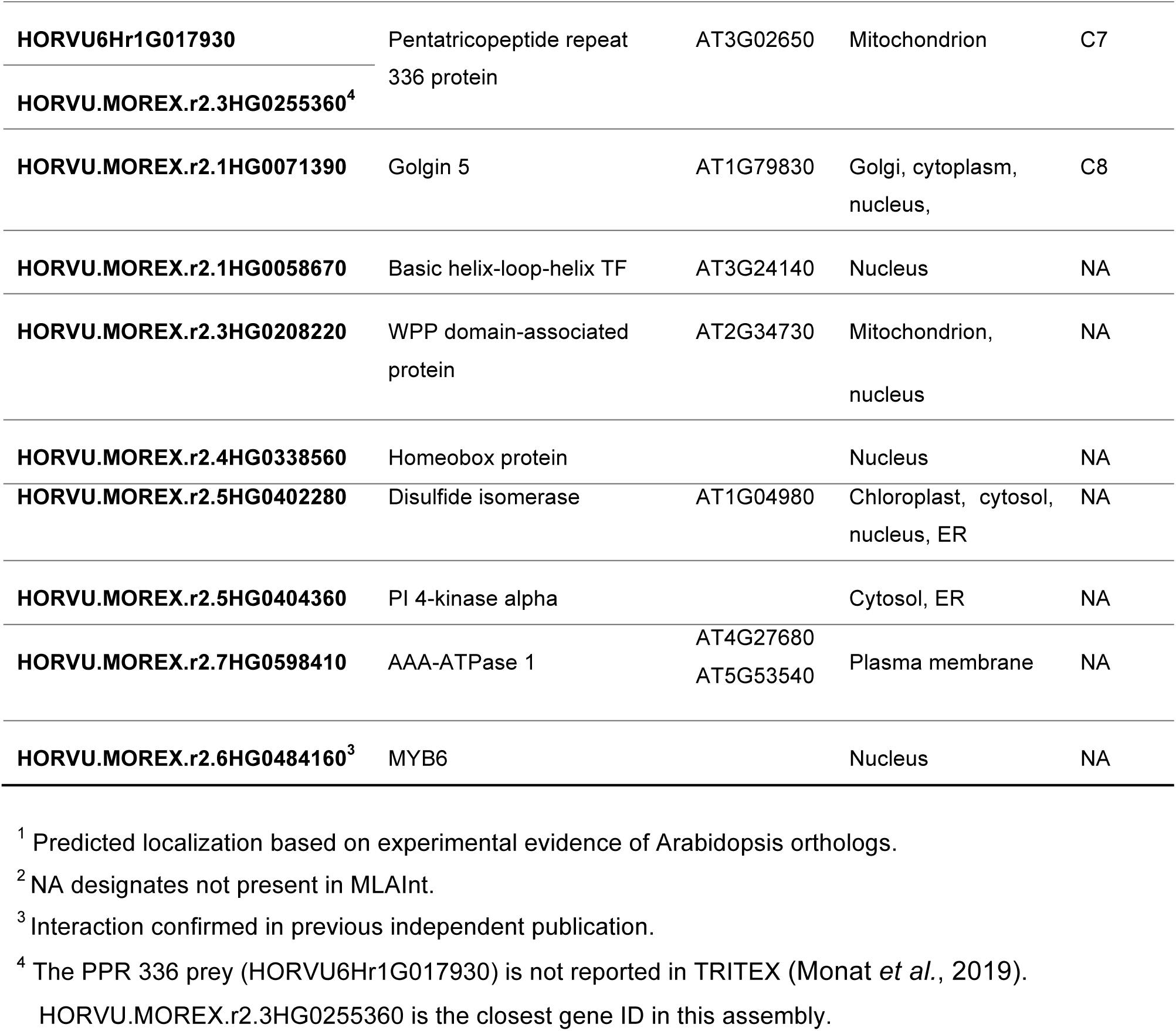
MLA_CC+NB_ validated interactors. Information including gene ID, description, *A. thaliana* ortholog, TAIR predicted cellular localization, presence in MLAInt and cluster number.

The CC+NB domains are highly conserved across MLA variants with more than 98% protein similarity (Seeholzer *et al*., 2010), therefore, we hypothesized that the validated interactors might be shared among multiple MLAs. We tested this by performing Y2H binary tests with the MLA13-type CC+NB domain present in MLA7, MLA9, MLA12 and MLA13, which differs from MLA6 by Q91K, P150T, and D177G amino-acid substitutions. MLA3 has a hybrid structure – the same as MLA13 at Q91K and P150T, but MLA6 at D177. MLA10 differs from MLA13-type by a single E41D change in the CC domain, and MLA1 and MLA8 is further diverged from MLA6, with S92F, L102F and P150T (Figure 4A) (Halterman and Wise, 2004; Seeholzer *et al*., 2010). All bait and prey sequences are reported in S1 Text. Positive interactions were found for all the validated targets with both the MLA6*-*type and MLA13*-*type CC+NB baits as shown in Figures 4B and S2, with some variation in the strength of the interactions for the two allele types. This suggests that these interactors are MLA-associated and that they are conserved across this NLR family.

**Figure 4.**
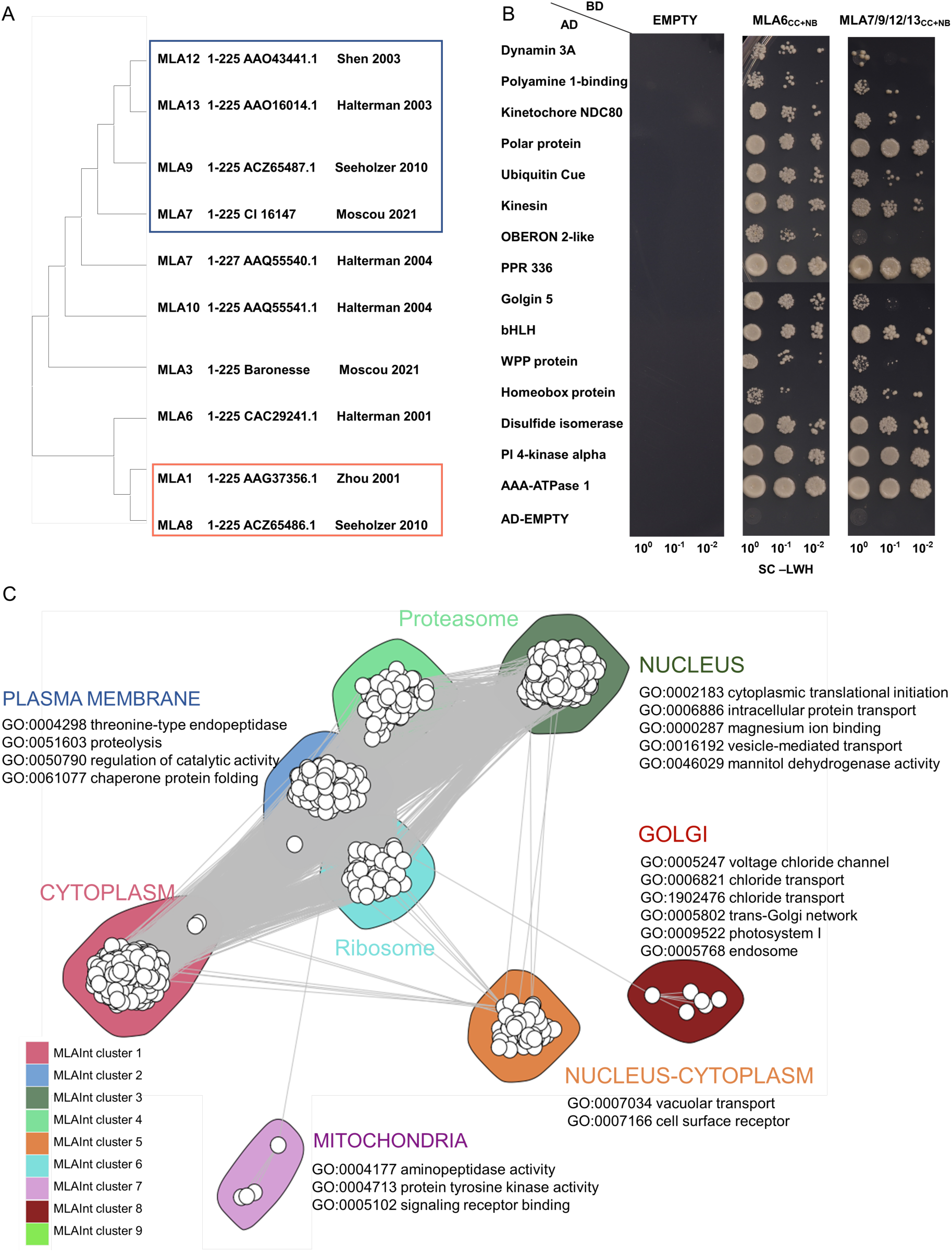
Novel interactors of MLA, conservation and signaling. A) Phylogeny of the amino acid CC+NB domains of *Mla* alleles, identical sequences are enclosed in boxes. B) Y2H binary confirmation of the interactors. Empty designates vector without insert. SC-LW media was used as control for diploid growth, and the interaction was tested using selection with SC-LWH media using three dilutions (10^0^, 10^−1^, 10^−2^). Order of preys is the same as in Table 1. C) Clustering of the MLA-associated interactome (MLAInt) with GO analysis.

We characterized two new MLA interacting TF, bHLH and HB (not present in HvInt), using phylogenetics and annotation of the TF gene families. Annotation included available knowledge of a role in immunity (Bruessow *et al*., 2021; Xu *et al*., 2014; Cabello *et al*., 2012; Gao *et al*., 2014; Coego *et al*., 2005), the presence of a Position Weight Matrix (PWM) (Weirauch *et al*., 2014; Matys *et al*., 2006; Jin *et al*., 2017), and the expression fold-change values between the wild-type CI 16151 (*Mla6, Bln1, Sgt1)* and its derived m18982 (*mla6, Bln1, Sgt1)* mutant, covering 0 to 48 HAI. We found 173 members of the bHLH family (Jones and Dangl, 2006) in barley, as shown in S3 Figure. The MLA-interacting bHLH interactor is divergent from other TFs characterized in defense response or possessing a PWM. The MLA-interacting bHLH appears to have a similar expression pattern in CI 16151 (*Mla6, Bln1, Sgt1)* and the susceptible mutant m18982 (*mla6, Bln1, Sgt1)* across the time course. We performed a similar analysis for the HB protein finding 109 family members. The MLA-interacting HB belongs to a previously studied clade, with a PWM reported and one related TF involved in plant resistance (Gao *et al*., 2014). The expression heatmap shows overexpression of the MLA-interacting HB in CI 16151 (*Mla6, Bln1, Sgt1)* as compared to m18982 (*mla6, Bln1, Sgt1)* (S3 Figure).

To explore signaling events triggered by MLA, we also positioned previously reported MLA protein interactors in the HvInt network (Table 1). These included HSP90, SGT1, MIR1, WRKY1, WRKY2, and MYB6 (Shen *et al*., 2007; Chang *et al*., 2013; Bieri *et al*., 2004; Wang *et al*., 2016). In total, thirteen targets of MLA were found in HvInt including eight validated in the current report using Y2H (Dreze *et al*., 2010), and five previously reported in the literature. We then computed an MLA-associated interactome (MLAInt), which consisted of proteins in the shortest paths between each pair of MLA interactors that do not pass-through MLA, adding the first and second neighbors to the resulting network (S5 Data). MLAInt has 1566 nodes and 13203 edges clustered into eight groups, with the thirteen MLA targets distributed in five of the MLAInt clusters (Figure 4C, Table 1). GO term analysis of the clusters points to different cellular processes associated with MLA-target signaling including vesicle-mediated transport (cluster 2), MAP kinase cascades (cluster 3), vacuolar transport, endosome, transmembrane receptor protein serine/threonine kinase activity (cluster 5), signaling and innate immune response (cluster 7), and trans-Golgi network and endosome (cluster 8).

The MLAInt subnetwork has significantly higher essentiality than HvInt in immunity (DiffSLC), degree centrality, betweenness and expression distance correlation, as supported by Wilcoxon rank sum tests (p-values 8.68×10^−93^, 0, 2.58×10^−29^, 1.04×10^−158^, respectively). The expression distance correlation was also compared by separating the datasets by defense phenotype for each subnetwork, finding higher values for MLAInt in all cases (Wilcoxon rank sum tests p-values 2.71×10^−53^ for resistant genotypes and 4.7×10^−33^ for susceptible genotypes). Then, we measured the distance path between MLA and the proteins in the HvInt, HvInt(R=S), and HvInt(R-S) networks. Distances between MLA and HvInt(R-S) are significantly lower than for HvInt or HvInt(R=S) (t-test p-values 2.4×10^−10^ and 1.2×10^−10^, respectively).

## DISCUSSION

In this report, we aimed to create a comprehensive overview of the different mechanisms that govern NLR-specified immunity in the large-genome cereal, barley (Deng *et al*., 2020). We used interolog inference to develop a predicted interactome (HvInt), which was integrated with context-specific datasets to model the immune response of barley to the powdery mildew pathogen. To start, we measured general network properties of HvInt and compared it to the Arabidopsis experimental interactome (AtInt), which we obtained by collecting 11253 proteins and 73960 experimentally validated interactions in this organism (S1 Data). We found the networks have common properties including a power-law distribution (scale-free) and small-world. These properties increase the robustness of the networks, buffering perturbation effects and optimizing information transfer, conditions that are maintained after sub-setting the original graph (Albert and Jeong, 1999; Watts and Strogatz, 1998). We extended these principles for the subsequent analyses that we performed using the HvInt and subnetworks, integrating them with barley defense-associated datasets, including results from eQTL, infection-time-course transcriptome and Y2H PPI interactions.

### Integration of eQTL and interactome data predicts disease modules during *Bgh* penetration and haustorial development

Disease-modules (Barabási *et al*., 2011) were used to characterize immune signaling at two key infection stages of barley by *Bgh*. Using GO terms and DE enrichment, we separated predicted module functions into core and unique responses. Unique responses at *Bgh* penetration (16 HAI) were associated with transcription and regulation of reactive oxygen species (ROS). For example, WRKY2, a previously reported interactor of MLA, was shown to be DM DE at this timepoint. WRKY2 has been shown to regulate the expression of genes involved in PAMP-triggered defense (Shen *et al*., 2007) and to be associated with race-non-specific resistance mediated by MLO (Spies *et al*., 2012). Other responses were associated with the regulation of ROS, including mitochondrial import inner membrane translocase subunit *Tim16/Pam16*, respiratory burst oxidase homolog (*RBOH*), and the N-carbamoyl-putrescine amidase. *Tim16/Pam16* and *RBOH* were overexpressed in the *mla6-*m18982 mutant as compared its progenitor, CI 16151 (*Mla6)*, suggesting that MLA has a regulatory role in stabilizing ROS functions. Indeed, the homolog of *Pam16* in *Arabidopsis* protects cells from over-accumulation of ROS species (Huang *et al*., 2013). This function may counteract the oxidative stress induced as a defense response associated with *Bgh* penetration (Trujillo *et al*., 2006). The N-carbamoyl-putrescine amidase is over-expressed in the CI 16151 progenitor at 16 HAI as compared to *mla6-*m18982. This enzyme participates in the biosynthesis of putrescine, a polyamine that induces ROS-dependent SA defense response and regulates *Bgh* penetration in the context of MLO resistance (Liu *et al*., 2020; Wu *et al*., 2017).

Transitioning to *Bgh* haustorial development (32 HAI), associated responses included phytoalexin biosynthesis, lytic enzymes, and pyruvate metabolism for a total of 67 DM DE genes. Phytoalexins participate in the SA immune response, a characteristic pathway against biotrophic pathogens (Dao *et al*., 2011). Not unexpectedly, we found chalcone synthase, a gene involved in phytoalexin metabolism, to be DM DE at 32 HAI. Chitinases, which we also found overexpressed at this timepoint, are secreted by the host cell to degrade fungal walls and elicit PAMP-triggered immunity (PTI) through chitin recognition (Pusztahelyi, 2018). Accumulation of chitinases has also been reported in the extra-haustorial complex during *Bgh* infection (Lambertucci *et al*., 2019). Similarly, other protein families that were also reported in the extra-haustorial complex were also found as DM DE at 32 HAI including genes associated with the TCA cycle and pyruvate metabolism, germin-like protein, and cysteine synthase. Genes involved in pyruvate metabolism are upregulated during haustorial development to contribute to energy production in defense and provide with precursors of other signaling pathways (Qiu *et al*., 2020; Lambertucci *et al*., 2019). Remarkably, among this group there is also a malate dehydrogenase (HORVU.MOREX.r2.1HG0067490) that was characterized as target of the *Bgh* effector BEC1054/CSEP0064 (Pennington *et al*., 2016).

We postulate that common GO terms at both stages of fungal development represent core responses that the host cell maintains during immunity, and these may be driven by the same or different genes. Often these core responses had annotations directly related to plant defense, including vacuolar transport, glutamate receptor (GLR) signaling and lignification. Vacuolar transport is a fundamental mechanism in plant response to pathogens, as multiple defense molecules are secreted to impair pathogen colonization (Yun and Kwon, 2017). DM DE genes associated with this function included KEULE and charged multivesicular body protein 5 (MVB5). Previous studies have found that the KEULE interacts with the SYP121 SNARE protein (ROR2), which confers penetration resistance of barley to *Bgh* and control the formation of the SNARE complex (Collins *et al*., 2003; Heese *et al*., 2001). MVBs have also been found to be transported to *Bgh* penetration sites and contribute to callose deposition (Böhlenius *et al*., 2010). We also found a GLR involved in potassium transport to be DM DE at both stages. This family of proteins has been found to act as sensors in plant resistance (Forde and Roberts, 2014). The *glr3*.*3* knock out mutant in Arabidopsis has shown increased susceptibility to obligate biotrophs such as *Hyaloperonospora* and modulation of ROS and nitric oxide (NO) production (Manzoor *et al*., 2013), while the triple mutant *glr2*.*7 2*.*8 2*.*9* is more susceptible to bacterial infection, dampening pattern-triggered immunity (PTI) signaling (Bjornson *et al*., 2021). Cinnamyl alcohol dehydrogenase was also found as DM DE in both timepoints, an enzyme that is involved in the lignification of the cell wall and in the activation of the salicylic acid (SA) defense pathway (Stadnik and Buchenauer, 2000). The above examples illustrate the power of the disease module concept, yet many more DM DE remain to be characterized during disease development (S3 Data).

### Resistant co-expression networks are associated with *Mla*

We used co-expression data to identify PPI subnetworks associated with disease phenotypes (Dong *et al*., 2015; Jiang *et al*., 2016; Mishra *et al*., 2017). Using distance correlation, we measured the non-monotonic behavior of our expression dataset (Székely *et al*., 2007) to obtain the resistant HvInt(R), susceptible HvInt(S) and the differentially co-expressed interactomes HvInt(R-S) and HvInt(S-R). The biological significance of these subnetworks lays on the assumption that co-expressed proteins are part of the same or overlapping pathways and coordinated by a common set of transcriptional regulators. At the physical level, the interactions in these networks may be part of the same protein complexes and then, high co-expression values increase the confidence of the predicted interactions. Our analyses indicate that the differentially co-expressed subnetworks, HvInt(R-S) and HvInt(S-R), have properties that predict their importance and role in defining the barley response to *Bgh*. For example, higher essentiality (DiffSLC), degree centrality and betweenness, and lower distance correlation, provide more robustness in signaling and carry more information (Koschützki and Schreiber, 2008; Mistry *et al*., 2017). Lower distance correlation values indicate a higher variability in the gene expression of the nodes, which is consistent with a differential response during defense between resistant and susceptible genotypes. Yet, despite the topological similarities between HvInt(R-S) and HvInt(S-R), only the resistant networks showed significant enrichment of eQTL associations with *Mla*.

GO enrichment of HvInt(R-S) indicates that this network is associated with multiple biological processes in plant defense, such as MAP kinase activity (Cui *et al*., 2019). Other GO terms including protein binding, folding, and ubiquitination were also enriched. The chaperone DnaK (*Hsp70*) and calreticulin were associated with these terms. Chaperones stabilize protein complexes and interactions. One *Hsp70* isoform has already been shown to participate in the response to salt, drought and heavy metal stress in barley, while another isoform is associated with response to fungal attack (Landi *et al*., 2019). Calreticulin is involved in calcium signaling and folding of glycoproteins, proteins from this family have been involved in the defense response to biotrophic pathogens, and to the stabilization of the EFR NLR in *Arabidopsis* (Qiu *et al*., 2012). Lastly, protein degradation via ubiquitination/proteasome 26S controls the dynamics of cellular functions such as hormone signaling, transcription and NLR-triggered immune response, including accumulation of MLA (Wang *et al*., 2016).

The enrichment of *Mla* eQTL associations in HvInt(R-S) indicates that these genes’ expression is controlled by signaling and transcriptional pathways in which *Mla* participates. Support for this hypothesis can be found when we consider that HvInt(R-S) was built from RNA-Seq data with active *Mla6* transcript accumulation [CI 16151 (*Mla6, Bln1, Sgt1)* and m19089 (*Mla6, bln1, Sgt1)*]. HvInt(S-R) in contrast was generated from mutants with active and inactive *Mla6* expression [m18982 (*mla6, Bln1, Sgt1)*, m11526 (*Mla6, Bln1, Sgt1*_*ΔKL308-309*_), and m19028 (*mla6, bln1, Sgt1)*] and had no enrichment of *Mla* eQTL associations. Further characterization of the predicted barley interactome can explain the association between MLA and proteins that regulate the expression of these genes. Incompleteness of the HvInt network limits this approach, as no interologs were found for most of the transcription factors that interact physically with MLA. Inference of gene regulatory networks can then be used to model the regulation mechanisms associated with MLA and to complement the analyses that we performed using PPI networks.

### MLA interactors confer information about receptor functionalities and its cellular localizations

Using Y2H-NGIS to mine interactions for different MLA6 domains resulted in differences in the number of positive validations. The highest number was obtained with the CC+NB fragment, domains that contain high protein similarity across MLA alleles. The CC domain has been described as one of the most abundant motifs in eukaryotes. It has been found to mediate specific interactions among multiple proteins involved in processes as diverse as cellular trafficking, assembly of complexes and transcription (Wang *et al*., 2012). Based on the functional evidence that links MLA_CC+NB_ self-association and activation of cell death, the reported interactors represent an important part of the MLA-immune signaling response. Previous studies have shown that MLA10_CC+NB_ self-interacts in yeast while shorter CC domains failed to reconstruct the associations, suggesting that MLA10_CC+NB_ has a higher stability (Maekawa *et al*., 2011). In addition, self-association of the CC domain is essential for immune signaling of the *Mla* orthologs Sr33 and Sr50 in wheat and rye, respectively (Casey *et al*., 2016) while triggering cell death in wheat (Cesari *et al*., 2016). At the structural level, available data shows different polymerization states across MLA and its orthologs. MLA10_CC_ appeared as a dimer in its crystal structure while Sr33_CC_ appeared as a tetramer (Casey *et al*., 2016; Maekawa *et al*., 2011). Higher order polymerization states have been reported for other NLRs, such as the pentamer obtained from the full length ZAR1 protein (Wang *et al*., 2019). Our interaction data suggest that the functionality of the CC domain may be linked to the NB domain, therefore it is possible that the polymerization of MLA reaches higher levels if a longer sequence of the protein is considered. Finally, multiple studies have shown that the N-terminal region of multiple NLRs is involved in triggering cell death. The domain of MLA10_CC_ activates cell death in *N. benthamiana*, and mutations of this domain impair the response (Maekawa *et al*., 2011).

By positioning the MLA interactors in HvInt, and subsequent subsetting to build MLAInt, we obtained insights about the function of the receptor. Using ortholog-aided association to its protein targets, we obtained a predictive model of MLA cellular localization during *Bgh* infection. Studies on MLA10 showed that cell death is triggered while the protein is localized to the cytosol, while disease resistance requires nuclear localization (Bai *et al*., 2012; Cesari, 2018; Shen *et al*., 2007). Our localization model indicates that one likely localization of MLA during the infection time course is the nucleus. This result is supported by previous evidence that points to an early nuclear localization of MLA during the *Bgh* infection time course (Shen *et al*., 2007). Our model suggests that MLA could be localized to the cytosol and possibly to other compartments as well, including the plasma membrane, the Golgi apparatus, the ER, and mitochondria. Here, we report a novel MLA interactor that may mediate the translocation of the MLA receptor between the nucleus and the cytoplasm, the polyamine-modulated factor 1-binding protein. The *Arabidopsis* ortholog of this protein, AT2G17990, is a calcium-dependent kinase adaptor protein involved in vacuolar biogenesis and trafficking (Kwon *et al*., 2018). This protein has one interactor reported in HvInt, the nuclear pore complex protein MOS7/NUP88 (Park *et al*., 2014), which is a constituent of a nuclear pore involved in the translocation and nuclear concentration of the R-protein SNC1, and the immune regulators EDS1 and NPR1 (Wiermer *et al*., 2010). This link between MLA and MOS7/NUP88 may indicate this nuclear pore mediates the translocation of the receptor to the nucleus.

Clusters of MLAInt coincide with different cellular locations. MLAInt cluster 3 has the highest number of MLA interactors with eight proteins predicted to be localized to the nucleus, including previously reported TFs WRKY1/2 and MYB6 (Chang *et al*., 2013; Shen *et al*., 2007). We report two novel MLA TF interactors including a bHLH and a HB protein, which were characterized using phylogenetic analysis as they were not found in the current HvInt. Results indicate that the MLA-interacting bHLH occupies an unexplored clade, separated from bHLH84, a TF that enhances autoimmunity of the NLR mutant *snc1* (Xu *et al*., 2014), and from bHLH059, a temperature-responsive salicylic acid immunity regulator in in Arabidopsis. The MLA-interacting HB protein was found to be closely related to the barley ortholog of *AtHB13*, a TF involved in tolerance to cold stress (Cabello *et al*., 2012) and further showed to confer resistance to powdery mildew, downy mildew, and green peach aphid in *Arabidopsis* (Gao *et al*., 2014). The clade that contains the MLA-interacting HB and the barley ortholog of *AtHB13* is separated from *OCP3*, another HB protein that confers resistance to necrotrophic pathogens in *Arabidopsis*, suggesting that the MLA-interacting HB may have specificity in the response to biotrophs. MLAInt cluster 5, the second group that contained nuclear MLA interactors, contained the OBERON and kinesin proteins. Functional annotations of this cluster indicate these proteins may be involved in different signaling pathways than the rest of the nuclear MLA interactors, with enrichment in vesicle transport and cell surface receptor signaling. According to our predictions, these two MLA targets share a series of receptor-like kinase interactors, important drivers of defense response. In *Arabidopsis*, OBERON and kinesin are hubs targeted by pathogen effectors from *Pseudomonas syringae* and *Hyaloperonospora arabidopsidis* (Mukhtar *et al*., 2011).

Presently, only the nucleus and the cytosol are reported cellular localizations of the MLA immune receptor (Shen *et al*., 2007). Our localization model points to other structures as well, although these are based on evidence from other species, and thus, need further study. First, we found a group of targets that shared nuclear and other cellular localizations such as the cell periphery and the ER. The POLAR protein in *Arabidopsis* relocates the BIN2 protein and other GSK3-like kinases from the nucleus to the plasma membrane regulating their activity and attenuating MAPK signaling (Houbaert *et al*., 2018). This mechanism may also mediate the translocation of MLA to the plasma membrane, considering that the NLR ZAR1 is positioned to this location to activate the immune response (Wang *et al*., 2019). The MLA interactor dynamin 3A, also predicted to localize to the plasma membrane, belongs to a protein family that in *Arabidopsis* controls cell death after powdery mildew infection (Tang *et al*., 2006). The ER was also found to be associated with MLA-signaling since the interactor disulfide isomerase (PDI), is predicted to be localized to this compartment. PDI is an enzyme that catalyzes protein disulfide bonds, helping to the correct folding and aggregation of proteins at this compartment (Ray *et al*., 2003). PDIs also have a role in the response to abiotic and biotic stresses (Kayum *et al*., 2017). Finally, we found two MLA targets with unique predicted cellular localizations, different than those already discussed: the pentatricopeptide repeat protein, assigned to MLAInt cluster 7, localized to the mitochondrion, and golgin 5 (MLAInt cluster 8), localized to the Golgi apparatus. Fluorescent expression of MLA10 indicates a protein distribution similar to the morphology of the Golgi apparatus (Shen *et al*., 2007). Golgin 5, localized to this compartment, is involved in vesicle tethering and regulation of intra-organelle transport by interacting with the small GTPase Rab6 (Latijnhouwers *et al*., 2007; Muschalik and Munro, 2018).

Our model for MLA signaling suggests that there is a transcriptional network mediated by MLA, which is likely triggered at early timepoints of infection. This response leads to the accumulation of MLA itself at about 16 HAI (at *Bgh* penetration). Our model also suggests that the immune receptor has functions in the cytoplasm and other cellular compartments, where it contributes to cell death. Further questions remain in this model, including where and when the recognition of effectors occurs and its influence in the translocation of MLA to different cellular compartments. Topological properties of MLAInt indicate a cohesive response of the subnetwork in defense signaling. In addition, distances between MLA and the proteins in HvInt(R-S) were shorter than for any other subnetwork, and enrichment of *Mla* eQTL associations was only found with resistant co-expressed subnetworks. This could indicate a potential involvement of the resistance protein in the signaling and transcriptional regulation of the proteins in HvInt(R-S). These results support a correlation between *Mla trans* eQTLs, MLA signaling and gene co-expression during resistance to *Bgh*.

## METHODS

### Collection of infection time course RNA-Seq and differential expression analysis

An infection time course of the wild-type CI 16151 (*Mla6, Bln1, Sgt1*) and derived fast-neutron mutants *bln1*-m19089 (*Mla6, bln1, Sgt1*), *mla6*-m18982 (*mla6, Bln1, Sgt1), rar3-*m11526 (*Mla6, Bln1, Sgt1*_*ΔKL308-309*_), and (*mla6+bln1)*-m19028 (*mla6, bln1, Sgt1*) was used for RNA-Seq analysis [NCBI-GEO GSE101304 (Chapman *et al*., 2020; Hunt *et al*., 2019)]. CI 16151 contains the functional *Mla6 R* gene and is resistant. *Bln1* (*Blufensin1*) is a negative regulator of PTI signaling, whose silencing results in down-regulation of genes associated with basal defense (Meng *et al*., 2009, Xu *et al*., 2015). The resistant *bln1* mutant, m19089, exhibits enhanced basal defense. *Mla6* is deleted in the m18982 mutant, and thus, is susceptible. *Rar3* (*required for Mla6 resistance3*) is required for *Mla6*-mediated generation of H_2_O_2_ and the hypersensitive response (HR). This susceptible mutant consists of a Lys-Leu deletion in the SGS domain of SGT1, which interacts with NLR proteins (Chapman *et al*., 2020). The (*mla6+bln1*) double mutant is susceptible as it contains the same *Mla6* deletion as in m18982.

First leaves were inoculated with fresh conidiospores from *Bgh* isolate 5874 (*AVR*_*a1*_, *AVR*_*a3*_, *AVR*_*a6*_, *AVR*_*a12*_) (Caldo *et al*., 2004) and sampled from a split-plot design at 0, 16, 20, 24, 32, and 48 HAI (5 genotypes × 6 time points × 3 biological replications). Raw reads (NCBI-GEO accession GSE101304) were processed using Trimmomatic (Bolger *et al*., 2014) and Salmon (Patro *et al*., 2017), taking as references the barley TRITEX (Monat *et al*., 2019) and *Bgh* DH14 (Frantzeskakis *et al*., 2018; Spanu *et al*., 2010) annotations. Taxon-specific normalization to the raw counts was applied: Salmon raw count matrices were separated for barley and *Bgh* and size factors were calculated using median-of-ratios normalization (Anders and Huber, 2010), which were then combined to calculate the final normalized counts matrix (Klingenberg and Meinicke, 2017). Differentially expressed (DE) genes were identified using a DESeq2 (Love *et al*., 2014) model with read counts as response, and timepoint and genotype terms as explanatory variables. We adjusted the p-values controlling for multiple testing, using Benjamin and Hochberg methodology (Benjamini and Hochberg, 1995), calling DE genes with an adjusted p-value of <0.001.

### Interactome reconstruction

We constructed a barley predicted protein-protein interactome, HvInt, using interologs (Matthews *et al*., 2001; Nakajima *et al*., 2018; Geisler-Lee *et al*., 2007). Orthologs of *H. vulgare* with *A. thaliana, Z. mays* and *O. sativa* were obtained using the Plant Compara tables V96 from Ensembl Plants (Howe *et al*., 2020) and with *Saccharomyces cerevisiae* using InParanoid8 (Sonnhammer and Östlund, 2015). These lists were filtered using the high confidence similarity scores from Ensemble and the maximum score cutoff of 100% from InParanoid8. Experimentally validated PPIs for these organisms were mined from BioGRID 4.1.190 version (Stark, 2005), the Protein-Protein Interaction database for Maize (PPIM) (Zhu *et al*., 2016), the Predicted Rice Interactome Network (PRIN) database (Gu *et al*., 2011), the pan-plant protein interactome (McWhite *et al*., 2020) and literature review (Mukhtar *et al*., 2011; Weßling *et al*., 2014; Smakowska-Luzan *et al*., 2018; Trigg *et al*., 2017; Wierbowski *et al*., 2020; Shen *et al*., 2007; Chang *et al*., 2013; Bieri *et al*., 2004; Wang *et al*., 2016). For each species, all the interactions were updated to the most recent Ensembl genome annotation and then, the compiled list was converted to barley gene IDs using the high confidence orthologs. The final non-redundant interactome was obtained by removing duplicated interactions and tracking the species where they were originally reported.

The infection-time-course expression dataset was integrated with the predicted barley interactome adding information to the edges and the nodes. We first assigned the weight of the edges as the pairwise distance correlation calculated from the complete expression dataset, or subsets of it based on the infection phenotype. Second, we adapted the DiffSLC (Mistry *et al*., 2017) pipeline to use expression data from RNA-Seq and compute the essentiality of the proteins in the predicted barley interactome. We characterized the predicted network, using clustering and GO term enrichment (Yu *et al*., 2012; Csardi and Nepusz, 2006). Scale-free and small-world network properties were tested for the final predicted interactome (Csardi and Nepusz, 2006; Watts and Strogatz, 1998; Albert and Jeong, 1999). Network topological properties such as protein essentiality with DiffSLC (Mistry *et al*., 2017), degree, betweenness (Csardi and Nepusz, 2006) and distance correlation (Székely *et al*., 2007) were used to characterize the subnetworks obtained from HvInt.

### Disease module prediction using eQTL and interactome data

We used N2V-HC (Wang *et al*., 2020) to integrate HvInt with eQTL data collected during *Bgh* appressorial penetration (16 HAI) and haustorial development (32 HAI) (Surana *et al*., 2017). All genetic markers with significant eQTL associations were used as input, removing associations with adjusted p-values larger than 0.001. The SNP and eQTL matrices used as input to the software were built to match the chromosome markers with linked genes (SNP table) and eQTL associations (eQTL table). The resulting modules for each timepoint were analyzed using GO enrichment. Results were compared and classified as core response (common between the two timepoints) and unique to each infection stage. Core and unique GO terms for the disease modules were analyzed using DE enrichment between the resistant CI 16151 (*Mla6, Bln1, Sgt1)* and the susceptible m18982 (*mla6, Bln1, Sgt1)* genotypes. If a gene associated with the GO term was DE, then it was called DM DE. Results from these analyses were summarized with a schema using Biorender.com. Expression profiles of the DM DE genes associated with each group were plotted using R (RCoreTeam, 2013).

### Construction of resistant and susceptible interactomes

Resistant and susceptible barley interactomes were obtained using expression distance correlation (Székely *et al*., 2007). The infection-time-course RNA-Seq was separated by disease phenotype and used to generate conditional subnetworks consisting of significantly co-expressed interactions (Jiang *et al*., 2016). The co-expression significance threshold used to generate each subnetwork was obtained by building a null distribution of this parameter. The RNA-Seq count data, grouped by phenotype, was permuted for each timepoint leaving the replicates as a block. The resulting distribution built from 10,000 permutations was used to calculate empirical p-values and correlation thresholds for significance of the pairwise correlation values. Using a p-value of 0.05, we calculated the significance correlation threshold for the resistant and susceptible subnetworks. Edges with values below the correlation thresholds were removed from the network to obtain the resistant [HvInt(R)] and susceptible [HvInt(S)] interactomes.

The differentially co-expressed resistant [HvInt(R-S)] and susceptible [HvInt(S-R)] subnetworks were obtained by removing the common interactions between HvInt(R) and HvInt(S), [HvInt(R=S)], from these subnetworks, respectively. The differentially co-expressed interactomes were further characterized using topological properties. Significance of the differences in these properties were calculated using Wilcoxon rank sum tests. Hypergeometric tests were applied to look for enrichment of eQTL associations in the HvInt(R), HvInt(S), HvInt(R-S), HvInt(S-R) and HvInt(R=S) subnetworks. The resistant subnetworks HvInt(R) and HvInt(R-S) were clustered using a walktrap algorithm (Csardi and Nepusz, 2006), and clusters were analyzed using GO enrichment (Yu *et al*., 2012). Visualization was done using Cytoscape (Shannon *et al*., 2003) and the R package RCy3 (Gustavsen *et al*., 2019).

### MLA-associated interactome from validated Y2H-NGIS data

Three MLA6 fragments (MLA6 aa 1-161 for the CC domain, MLA6 aa 1-225 for CC and NB domains, and MLA6 aa 550-956 for the LRR domain) were tested as baits using Y2H-NGIS, and a three-frame cDNA prey library of 1.1 ×10^7^ primary clones generated from the infection time course experiment (Hunt *et al*., 2019; Surana, 2017; Velásquez-Zapata *et al*., 2021). Y2H-NGIS data from these baits were analyzed using the NGPINT and Y2H-SCORES pipelines (Banerjee *et al*., 2021; Velásquez-Zapata *et al*., 2021). Using the Borda ensemble provided by Y2H-SCORES, we predicted a top list of candidate interactors to be validated. Interacting prey fragments of the top interactors were determined using the IGV alignments obtained from the NGPINT and the reported *in-frame* prey transcripts with the highest *in-frame* score from Y2H-SCORES. Primers were designed for Gateway cloning, and clones were inserted into the prey vector. After transforming the candidates into yeast, we performed a binary Y2H test (Dreze *et al*., 2010) under three levels of selective media: Diploid selection (SC-LW), interaction selection (SC-LWH) and selection (SC-LWH) using three dilutions (10^0^, 10^−1^, 10^−2^). These interactions were extended to the CC+NB domain sequences of the MLA13-type alleles MLA7, MLA9, MLA12, and MLA13. All bait and prey sequences are reported in S1 Text.

The MLAInt subnetwork was built by retaining the nodes in the shortest paths computed for any pair of MLA interactors. First and second neighbors to the nodes in this subnetwork were added to complete the MLAInt. This subnetwork was clustered using a fast-greedy algorithm (Csardi and Nepusz, 2006), and clusters were analyzed using GO (Yu *et al*., 2012) and eQTL (Surana *et al*., 2017) enrichment. Topological properties were used to characterize MLAInt. Visualization was done using igraph (Csardi and Nepusz, 2006).

### Phylogenetic trees of predicted barley transcription factor families

A total of 173 and 109 predicted barley bHLH and homeobox transcription factors were predicted, using the TRITEX Morex r2 barley high confidence protein reference and PlantTFDB 4.0 (Monat *et al*., 2019; Jin *et al*., 2017). The sequences were aligned using MUSCLE (Edgar, 2004), and then the trees were calculated using maximum likelihood with a Jones-Taylor-Thornton substitution model and a gamma-distributed-rate among sites model. The resulting trees were annotated using the ggtree R package (Yu *et al*., 2017). The annotation consisted in plotting: 1) genes with PWM DBD domain according to our database search (Weirauch *et al*., 2014; Jin *et al*., 2017; Matys *et al*., 2006); 2) Log2 of the foldchange of the RNA-Seq expression data for the comparison between CI 16151 (*Mla6, Bln1, Sgt1)* wild-type genotype and the m18982 (*mla6, Bln1, Sgt1)* mutant in a time course experiment of infection (0, 16, 20, 24, 32, 48 HAI); and 3) previously characterized TFs in plant immunity.

## Availability of code, data, and materials

All the code to support the main findings of this manuscript can be found at the GitHub page https://github.com/Wiselab2/Barley_Interactome. All MIAME-compliant Barley1 GeneChip profiling data (Affymetrix part number 900515) (Close *et al*., 2004) from the Q21861 x SM89010 eQTL analysis are available as accession GSE68963 at NCBI’s Gene Expression Omnibus (GEO) (Surana *et al*., 2017). Conversion of the GeneChip Probe sets to gene IDs was done using Ensemble and Biomart (Howe *et al*., 2020), Infection-time-course RNA-Seq datasets are available in NCBI-GEO under the accession number GSE101304 (https://www.ncbi.nlm.nih.gov/gds/?term=GSE101304). R code and the ReadMe file for the NGPINT and Y2H-SCORES software used to identify MLA interactors are provided at the GitHub page https://github.com/Wiselab2/. Raw Y2H-Seq reads are at NCBI-GEO under accession numbers GSE164814 (MLA6_1-161_), GSE164815 (MLA6_1-225_), GSE164816 (MLA6_550-956_), GSE164761 (luciferase - construct #1) and GSE164762 (luciferase - construct #2).

## Abbreviations

AtInt: *Arabidopsis thaliana* interactome
*Bgh*: *Blumeria graminis* f. sp. *hordei*
bHLH: basic Helix-Loop-Helix
CC: Coiled-Coil
DE: Differentially Expressed
DM: Disease module
eQTL: expression Quantitative Trait Loci
ER: Endoplasmic Reticulum
GLR: Glutamate Receptor
GO: Gene Ontology
HAI: Hours After Inoculation
HB: Homeobox
*Hv*: *Hordeum vulgare*
HvInt: *Hordeum vulgare* predicted interactome
LRR: Leucine-Rich Repeat
*Mla1*: powdery mildew resistance locus a1
MVB5: charged Multivesicular Body protein 5
NBS: Nucleotide-Binding Site
NO: Nitric oxide
NLR: Nucleotide-binding leucine-rich repeat
PPI: Protein-Protein Interaction
RBOH: Respiratory Burst Oxidase Homolog
ROS: Reactive oxygen species
SA: Salicylic Acid
Y2H: Yeast-Two-Hybrid
Y2H-NGIS: Yeast-Two-Hybrid, Next-Generation Interaction Screening

## Supplemental files

S1 Figure. Core and unique DM GO terms associated with DE genes by timepoint.

S2 Figure. Y2H validation for the CC+NB domains of *Mla* alleles.

S3 Figure. Phylogenetic analysis of the bHLH and HB families in barley.

S1 Data. Predicted barley interactome (HvInt) and experimentally validated Arabidopsis interactome (AtInt).

S2 Data. Node properties of HvInt and enrichment of essential proteins across clusters.

S3 Data. Disease modules and DM DE genes at *Bgh* penetration and haustorial development.

S4 Data. Resistant and susceptible interactomes. S5 Data. MLA interactome.

S1 Text. Bait and prey sequences used for Y2H binary validation among *Mla* alleles.

## Acknowledgments

The authors thank Sagnik Banerjee for his invaluable assistance mapping of the RNA-Seq raw reads to the TRITEX genome, and Ana Mía Corujo Ramirez, Morgan Bixby, Jessica Faust, and Stephanie Schuler for technical assistance with the Y2H-NGIS screens. We thank Matt Moscou for sharing sequences corresponding to the CC+NB of *Mla3* and *Mla7*. Research supported in part by Fulbright - Minciencias 2015 & Schlumberger Faculty for the Future fellowships to VVZ, USDA-ARS Postdoctoral Research Associateship and USDA-NIFA-ELI Postdoctoral Fellowship 2017-67012-26086 to JME, and National Science Foundation - Plant Genome Research Program grant 13-39348, USDA-National Institute of Food and Agriculture grant 2020-67013-31184 and USDA-Agricultural Research Service project 3625-21000-067-00D to RPW. The funders had no role in study design, data collection and analysis, decision to publish, or preparation of the manuscript. Mention of trade names or commercial products in this publication is solely for the purpose of providing specific information and does not imply recommendation or endorsement by the USDA, NIFA, ARS, or the National Science Foundation. USDA is an equal opportunity provider and employer.

## Author Contributions

Designed the research: VVZ, JME, RPW.

Performed research: VVZ, JME, GF.

Contributed new analytic/computational tools: VVZ.

Analyzed data: VVZ.

Wrote the paper: VVZ.

Edited the paper: VVZ, GF, RPW.

